# Trehalose Stabilizing Protein in a Water Replacement Scenario: Insights from Molecular Dynamics Simulation

**DOI:** 10.1101/2019.12.27.889063

**Authors:** Qiang Shao, Jinan Wang, Weiliang Zhu

## Abstract

How trehalose has exceptional property in helping biomolecules preserve their native structures remains a subject of active research. Running molecular dynamics simulations on a model protein in low-concentrated trehalose solution and pure water, respectively, the present study verifies the ability of trehalose in stabilizing protein native structure and provides a comprehensive atomic-level picture of the molecular interactions among protein, trehalose, and water in their mixed solution. Trehalose directly interacts to and meanwhile affects the interactions between the other species *via* hydrogen bonding: 1) trehalose molecules are clustered through inter-molecular hydrogen bonding interaction; 2) trehalose forms hydrogen bond with water which influences the strength of water-water hydrogen bonding network but does not impair protein-water hydrogen bonding; 3) trehalose is accessible to form hydrogen bonds towards protein and simultaneously replace water molecules around protein which reduces the hydrogen bonding possibility from water to protein, in accordance with “water replacement” scenario.

## Introduction

It is well recognized that saccharides have exceptional properties in helping biomolecules such as proteins preserve their native structures under harsh conditions, e.g., high or low temperatures, and dehydration [1,2]. Among naturally available saccharides, trehalose, a nonreducing homo-disaccharide in which two D-glucopyranose units are linked together by an α-1,1-glycosidic linkage (Figure 1 (a)) is probably the best biomolecule stabilizer [3–8]. As a result, trehalose is widely used as additive for long-term preservation of therapeutic proteins, foods, and cosmetics industries [9–11].

**Figure 1.**
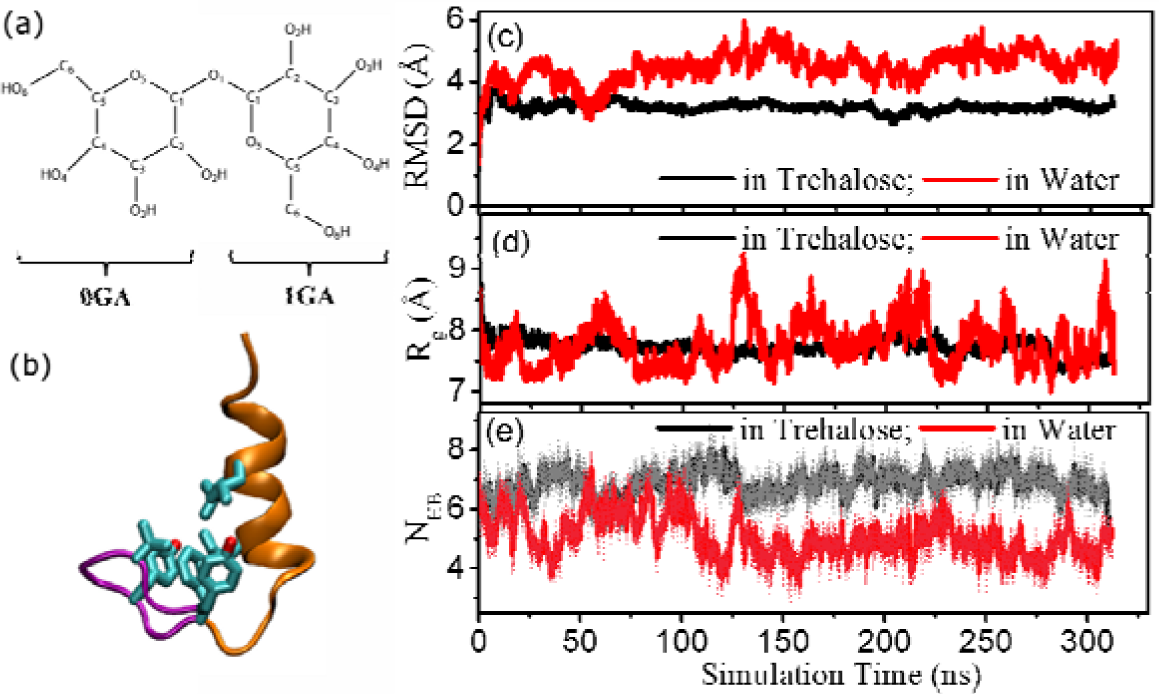
(a-b) Structure of trehalose molecule and BBA5 protein, respectively. (c-e) Time series of the root-mean-square deviation (RMSD) corresponding to the NMR structure, the radius of gyration (R_g_), and the total number of backbone hydrogen bonds formed within BBA5 protein in trehalose solution and pure water, respectively.

A great amount of experimental and theoretical studies have been performed to investigate the dynamic properties of trehalose, the molecular interactions among trehalose, water, and proteins in their mixed solution, with the hope to reveal the molecular mechanism underlying the prominent stabilizing effects of trehalose towards proteins [3,6,12–28]. Multiple hypotheses have been proposed accordingly. Mechanical entrapment (vitrification) hypothesis suggests that trehalose molecules form a highly viscous glassy matrix, likely vitrifying the solution which causes the motional inhibition of proteins and thus kinetically maintains their structures [18,19]. Water replacement hypothesis, on the other hand, proposes that trehalose could replace (most of) water molecules in the first solvation shell of protein and in the meanwhile form direct hydrogen bonds towards protein, which satisfy the hydrogen bonding requirement from protein surface polar groups, preserving the native structure of protein [3,20,21]. Water entrapment hypothesis, however, emphasizes that protein structure maintenance should be mainly attributed to the hydration water molecules trapped in the intermediate layer between trehalose and protein surface, implying that the glassy matrix constructed by trehalose is capable of concentrating residual water molecules around protein [22]. In addition, the observation by elastic neutron scattering and Raman scattering experiments [23–25] on binary sugar-water systems that trehalose alters the surrounding hydrogen bonding network of water solvent suggests that trehalose could exert an indirect influence on the dynamics of protein molecules.

Each abovementioned model is one way or another supported by a variety of experiments and/or theoretical simulations. For instance, extensive research work on proteins embedded in amorphous trehalose matrixes have evidenced strong dynamical coupling between protein and surrounding trehalose matrix, providing valuable information on trehalose controlled protein dynamics (mechanical entrapment hypothesis) in solutions consisting of very high-concentrated trehalose and low water content [29–36]. Whether the mechanical entrapment hypothesis can be still used to explain the good protein-stabilizing properties of trehalose in low and medium concentrated solutions (such as in natural condition in living organisms) is, however, questionable. As a matter of fact, the Fourier transform infrared spectroscopy experiment did show that the formation of amorphous phase alone in not sufficient to maintain protein structure during dehydration [37]. This experiment along with others pointed out that both the formation of a glassy trehalose matrix and the hydrogen bonding from trehalose towards protein play important roles in maintaining protein structure although the relative contribution of the two factors is uncertain [3,37–39]. Meanwhile, there are also many experimental and theoretical studies indicating that the interactions from trehalose and the induced effects on protein are better described in terms of water entrapment hypotheses [17,36,40–42]. Therefore, no generally accepted answer has been provided yet for the question why trehalose has superior protecting function towards biomolecules, making it remain a subject of active research.

In view of this, to provide a comprehensive atomic-level picture of the dynamical properties of trehalose and its interactions with protein and water in protein-trehalose-water complex solution, the present study ran extensive molecular dynamics (MD) simulations on a *de nono* designed *ββα* mini-protein BBA5 (PDB code: 1T8J [43], see its structure in Figure 1 (b)) dissolved in both 0.6 M trehalose aqueous solution and pure water. Three independent trajectories were run for each simulation system to guarantee the convergence of computational calculation. The comparative study demonstrates apparent stabilizing function of trehalose towards protein even in low-concentrated trehalose/water mixture, consistent with previous many research studies [12,13,44]. The detailed analysis of interactions among protein, trehalose, and water from structural and energetic perspectives ascertains the specific interactions which play crucial role in affecting the dynamics of protein, suggesting that the protein stabilization in (low-concentrated) trehalose solution might mainly follow the water replacement hypothesis.

## Materials and Methods

All MD simulations were performed in explicit solvent at room temperature, making use of AMBER 11 suite of programs [45] with FF99SB molecular force field [46]. Trehalose molecule is modeled by using GLYCAM06 force field [47] and water is described with TIP3P explicit solvent model [48]. Three independent trajectories were run to test the calculation convergence for each simulation system. Detailed parameters of individual trajectories are presented in Table S1 in Supplementary Materials.

For BBA5 in 0.6 M trehalose solution, the native structure of BBA5 (PDB code: 1T8J [43]) was immersed into a cubic box containing 120 trehalose and 6949 water molecules. The cubic box set for BBA5 in pure water contained 5639 water molecules. One Cl^−^ anion was then added in each system to balance the system charge. For each simulation system, the simulation procedure included the energy minimization, the following heating-up process, and the final long-time equilibrium simulation calculation (production run). NPT (number, pressure, temperature) ensemble calculations were performed and periodic boundary conditions were used in the simulations. The pressure of the system was set as 1 atm and the temperature was controlled as 300 K. The SHAKE algorithm [49] was used to constrain all bonds involving hydrogen atoms. A cutoff of 10.0 Å was applied for nonbonding interactions. The Particle Mesh Ewald method was applied to treat long-range electrostatic interactions [50]. The Langevin dynamics with a collision frequency of 3.0 ps^−1^ was adopted to control the temperature of the system.

The energy of each system was minimized through a total of 2500 steps of calculations: 1000 steps of steepest descent minimization with the polypeptide being fixed using harmonic restraints (using a force constant of 500.0 kcal mol^−1^ Å^−2^ to apply to the backbone atoms), which was then followed by 1500 steps of conjugate gradient minimization. Subsequently, to better relax the system, the system was heated to 360 K and equilibrated for several nanoseconds at 360 K, which was followed by a 1 ns of cooling from 360 to the target temperature 300 K. During the heating and cooling processes, the protein configuration kept fixed by applying harmonic restraints on the backbone atoms (force constant = 10.0 kcal mol^−1^ Å^−2^). The three independent trajectories of each solution system are different in the running time used to equilibrate the system at 360 K. Finally, longtime production runs were performed at 300 K until the protein reached its equilibrium structure (~330 ns per trajectory). The data in each trajectory was collected every 2.0 ps.

## Results

### Enhanced Stability of Protein Native Structure in Trehalose Solution

NMR experiment [43] displayed that BBA5 protein adopts a simple tertiary structure consisting of a β-hairpin (Tyr1-Phe8) segment and an α-helix (Arg10-Gly23) segment in water (Figure 1 (b)). Both segments are packed with each other through intra-protein hydrophobic interactions. Here, the structural stability of BBA5 was investigated in trehalose solution. In addition, the same protein was also investigated in pure water as a control test. For each simulation system, three independent simulation trajectories were run, each lasting ~330 ns. Within the simulation times, three trajectories reached similar (solution-dependent) stable conformations, indicating the convergence of simulation calculation. The data analysis in the present study (Figures 1 to 8 and Table 1) is based on the average of three trajectories, which we believe could provide more reliable information for protein motion in solution than single trajectory does. The single trajectories for BBA5 protein in trehalose or water solution are present in Figures S1-S2 in Supplementary Materials.

**Table 1.**
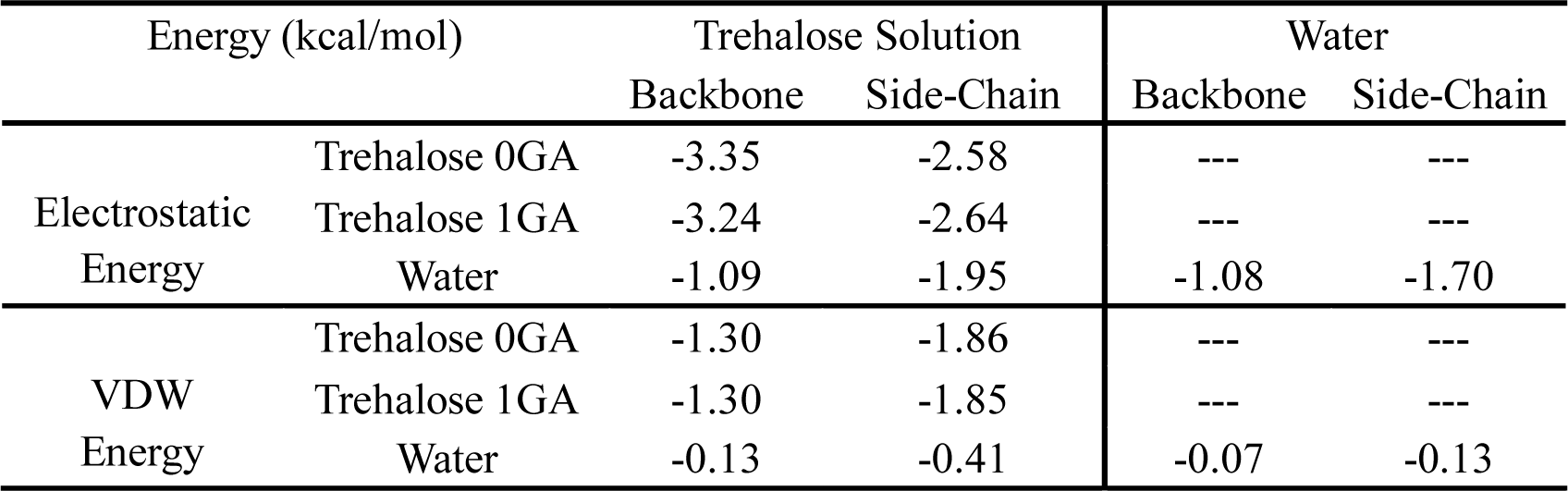
Decompositions of nonbonded energy between each trehalose/water and backbone and side-chain of protein.

| Energy (kcal/mol) |  | Trehalose Solution |  | Water |  |
| --- | --- | --- | --- | --- | --- |
|  |  | Backbone | Side-Chain | Backbone | Side-Chain |
| Electrostatic<br>Energy | Trehalose 0GA | -3.35 | -2.58 | --- | --- |
|  | Trehalose 1GA | -3.24 | -2.64 | --- | --- |
|  | Water | -1.09 | -1.95 | -1.08 | -1.70 |
| VDW<br>Energy | Trehalose 0GA | -1.30 | -1.86 | --- | --- |
|  | Trehalose 1GA | -1.30 | -1.85 | --- | --- |
|  | Water | -0.13 | -0.41 | -0.07 | -0.13 |

Figure 1 (c-d) illuminates the time evolution of the root-mean-square deviation (RMSD) and the radius gyration (Rg) of BBA5 in trehalose solution and pure water, respectively. Both RMSD and Rg values of the protein keep small and steady in the entire simulation of trehalose solution whereas the two types of parameters fluctuate in a large range in pure water. Accordingly, the root-mean-square fluctuations (RMSFs) of individual residues, which can be used to evaluate the structural flexibility of protein, also have much smaller values in trehalose solution than in pure water (Figure S3 in Supplementary Materials). In addition, the total number of backbone hydrogen bonds formed within BBA5 (N_HB_) in trehalose solution is greater than that in pure water (Figure 1 (e)), indicating the better maintenance of protein secondary structure in the former solution. Therefore, the presence of trehalose enhances the stability of the global native structure of BBA5.

To see more clearly the stabilizing effects of trehalose on the structure of BBA5, the solvent accessible surface area (SASA) of BBA5 and its polar and nonpolar components were calculated. As shown in Figure 2, the total SASA of BBA5 in trehalose solution fluctuates in a relatively narrower range than the counterpart in pure water does. More specifically, the polar SASA is slightly influenced but the nonploar SASA is more biased to the small value in the presence of trehalose, which leads to more rigid protein structure in trehalose solution.

**Figure 2.**
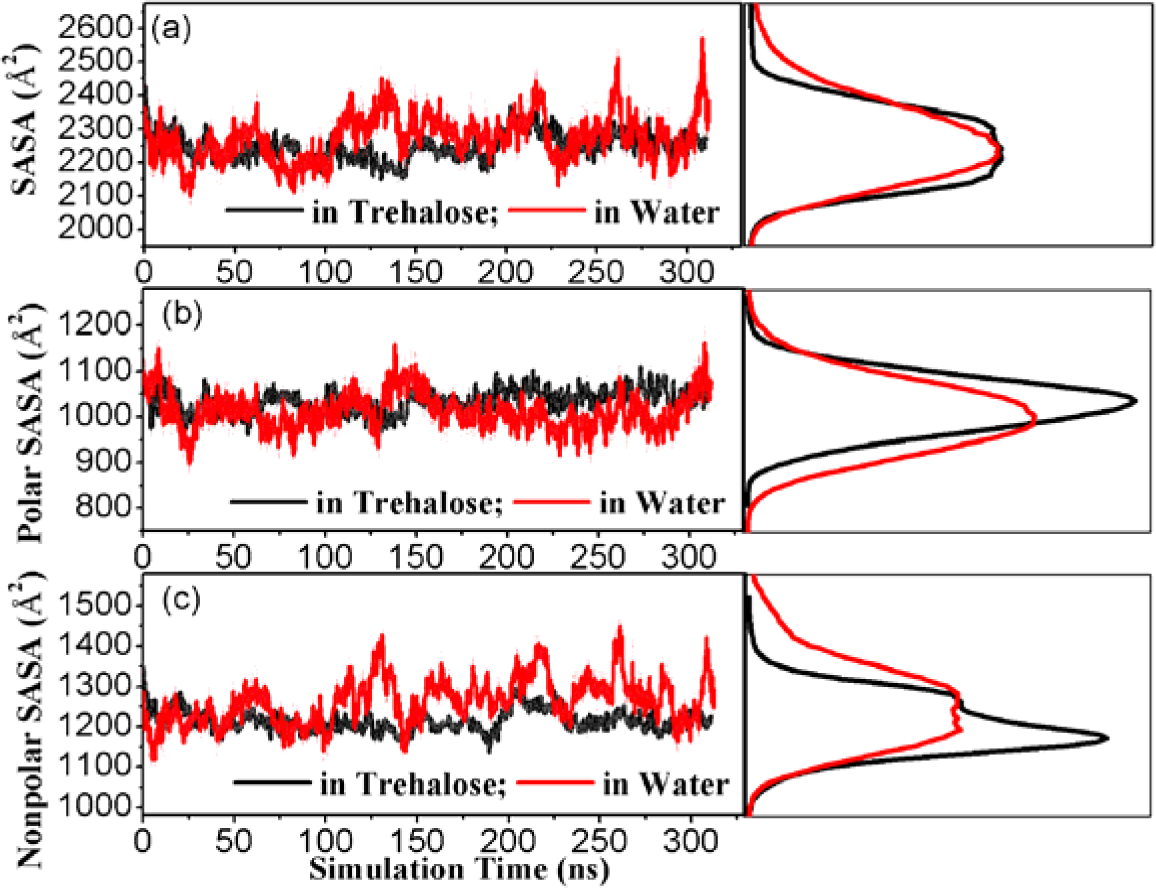
Time series of (a) the total solvent accessible surface area (SASA) of BBA5 protein, and (b) the polar and (c) nonpoar components, along with their respective distribution in trehalose solution and pure water.

### Water Molecule Expellation from Protein Surface by Trehalose

In “water replacement hypothesis” [3,20,21], trehalose is proposed to replace hydration water molecules around protein and in the meanwhile form direct hydrogen bonds towards protein. To see the distribution of trehalose and water molecules surrounding protein, we calculated the time series of the number of the two species in the first solvation shell (FSS) of protein (3.4 Å around protein surface) in trehalose solution.

One can see from Figure 3 that trehalose molecules have a tendency to approach protein surface along simulation, as revealed by the gradually increased number of trehalose in the FSS of protein. The approach of trehalose to protein suface is not a rapid motion, which is completed at around 150 ns and afterwards the trehalose number around protein keeps steady. Even so, the trehalose molecules in close proximity to protein only account for around 20% of the whole trehalose throughout the solution. In comparion to the molar ratio of trehalose to water in the whole solution system (17.3%), the trehalose:water ratio in the FSS of protein is slightly increased (~20.0%), indicating moderate tendency of trehalose clustering around protein. Nevertheless, as a direct consequence of the clustering of large-size trehalose, water molecules are highly expelled from the FSS of protein. In compaison with the large number of water molecules staying around protein in the simulation of pure water, about one third of water molecules are removed from protein surface in trehalose solution (Figure 3).

**Figure 3.**
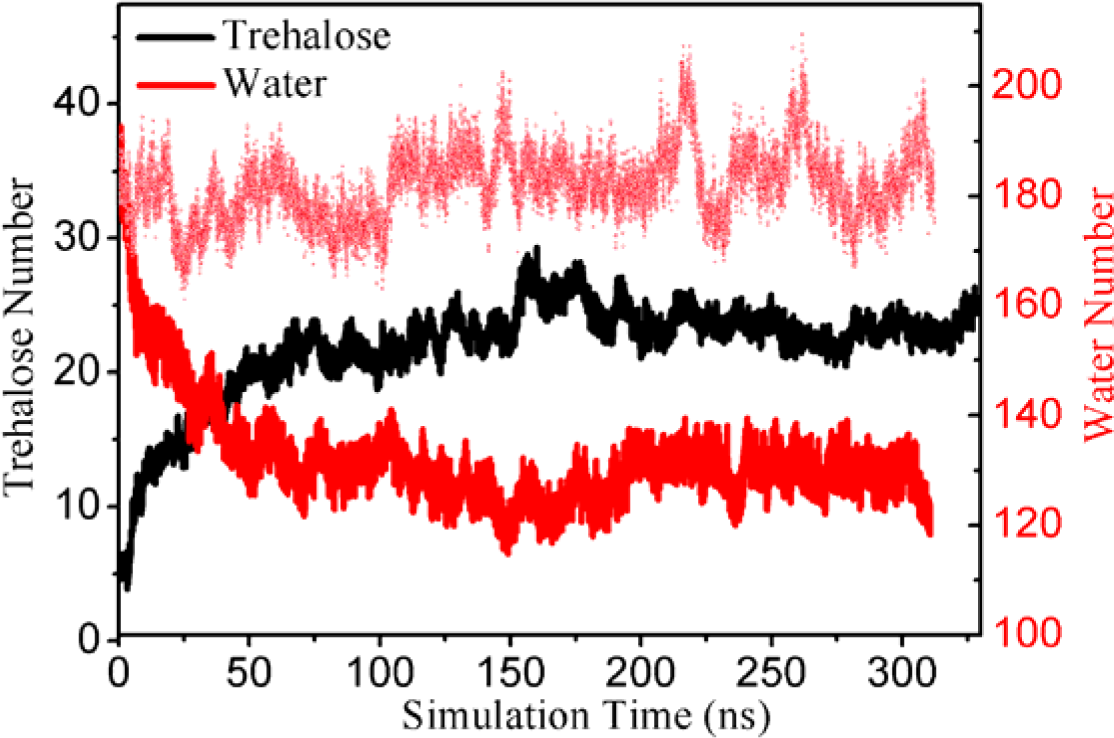
The number of trehalose and water molecules in the first solvation shell of BBA5 protein as a function of simulation time for BBA5 in trehalose solution. The number of water molecules in the first solvation shell of BBA5 in pure water is represented by dash red line to show different hydration level around protein surface in the two solutions under study.

The moderate increase in the fraction of trehalose in the FSS of protein relative to bulk solution can be explained from energetic perspective [51,52]. The electrostatic energy between each trehalose in proximity to protein (defined as the area within 5.0 Å of protein) and in bulk region (6.0 Å away from any protein atoms), respectively, with the rest of the system was calculated. The van der Waals (VDW) interaction energy of each water (and particularly trehalose) with the rest of the system is positive, which indicates unfavorable VDW interaction among water and trehalose in solution, is not presented. One can see from Figure 4 (a-b) that either 0GA or 1GA ring of trehalose has similar value of electrostatic energy, suggesting that the two parts contribute equally in the inter-molecular interactions of trehalose. In addition, the trehalose molecule around protein has a distribution of electrostatic energy with a broader peak at lower energy range than that in the bulk (energy difference is −7.76 kcal/mol). Consequently, it is the more favorable electrostatic interactions between trehalose and protein that drive trehalose molecule to protein surface. Interestingly, although the electrostatic energy difference between water in the region of protein surface and water in bulk solution is not changed in both trehalose solution and pure water, the detailed value of electrostatic energy per water (either around protein or in the bulk) is more negative in trehalose solution than that in pure water. Therefore, it is reasonable to speculate that the presence of trehalose somehow results in more favorable electrostatic interaction of water in aqueous solution.

**Figure 4.**
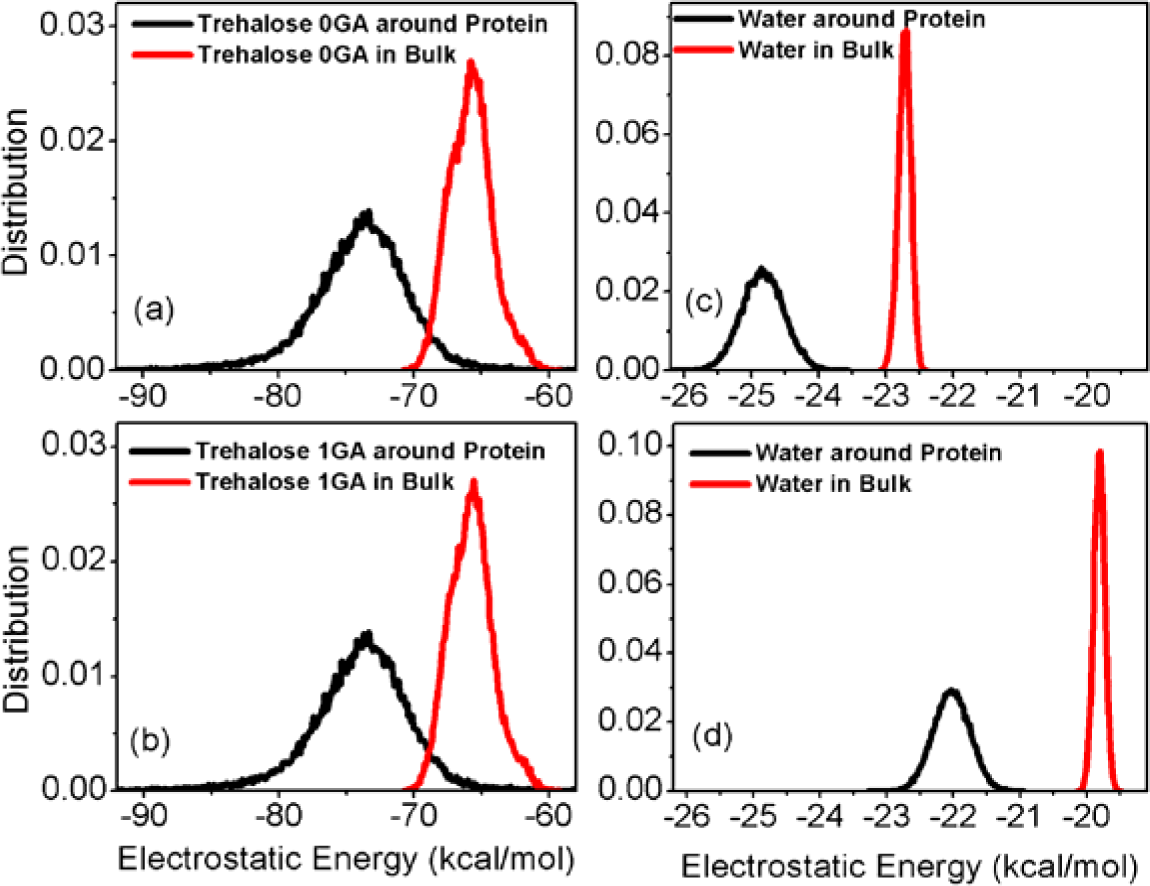
(a-b) Distribution of the electrostatic energy of 0GA and 1GA segments of each trehalose in proximity to protein and in the bulk region, respectively, with the rest of system for BBA5 in trehalose solution. (c) Distribution of the electrostatic energy of each water in proximity to protein and in the bulk region, respectively, with the rest of system for BBA5 in trehalose solution. (d) Distribution of the electrostatic energy of each water in proximity to protein and in the bulk region, respectively, with the rest of system for BBA5 in pure water.

The favorable protein-solvent (co-solvent) interactions can be further indicated by the detailed decomposition of interaction energies of single trehalose/water molecule (around protein surface) with protein only. As shown in Figure 5, both 0GA and 1GA rings of trehalose have similar negative electrostatic energies with protein, indicating trehalose has strong electrostatic interaction with protein. Meanwhile, trehalose could also have VDW interaction with protein, consistent with the proposal by Koa et al. that trehalose, as a moderate amphiphile, possesses both hydrophobic and hydrophilic moities [53]. Water molecule, on the other hand, can only have electrostatic interaction with protein, as revealed by its negative electrostatic energy but not VDW energy towards protein. In addition, the average electrostatic interaction per water to protein is more or less similar in trehalose solution and pure water, indicating that the presence of trehalose doesn’t impair the interaction strength for each water to protein. Table 1 lists the average electrostatic and VDW energies per trehalose/water molecule to protein and the decomposition into protein backbone and side-chain parts, respectively. Either the electrostatic and VDW energies of trehalose or the electrostatic energy of water more or less equally contributes to the backbone and side-chain of protein.

**Figure 5.**
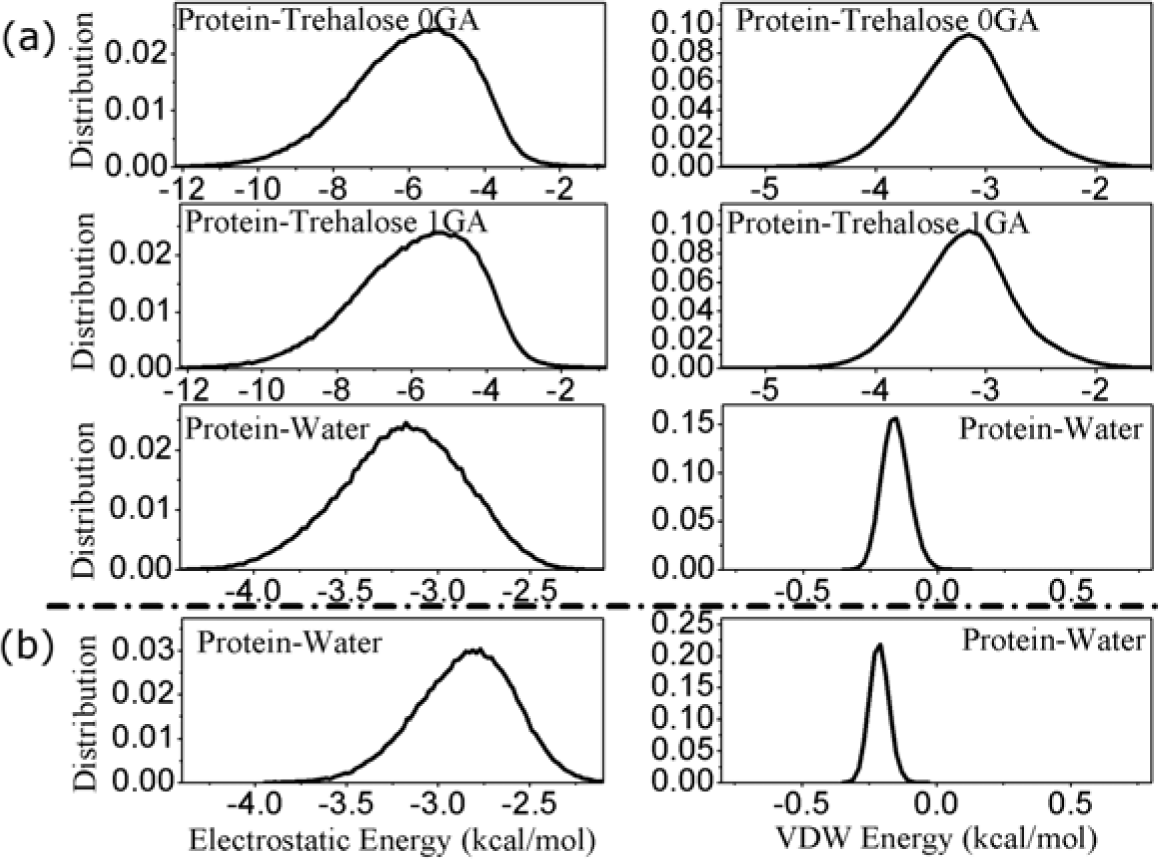
Distribution of electrostatic and VDW energies between protein and each trehalose/water (in proximity to protein) for BBA5 in (a) trehalose solution and (b) pure water.

### Undisturbed Protein-Water but Enhanced Water-Water Hydrogen Bonding Strenght in the Presence of Trehalose

The radial distribution function (RDF) can be used to describe the average structure of liquid. Figure S4 (a-b) in Supplementary Materials shows the RDF of different atoms of trehalose aound protein backbone carbonyl (-C=O) and amide (-NH) groups in trehalose solution. Considering the large number of atoms of individual trehalose molecule, here we used three representatives but not all oxygen atomes, namely, O_1_, O_5_, and O_6_ in the RDF calculation (the hydroxyl oxygens of O_2_, O_3_, and O_4_ behave in a similar fashion as that of O_6_ and thus are not presented). High peaks can be seen in the RDF diagrams of the O_6_ atom of trehalose around both backbone carbonyl and amide groups, demonstrating the formation of hydrogen bonds between the protein backbone and trehalose (mainly on the hydroxyl groups of O_6_ (and O_2_, O_3_, and O_4_)). The same shape of RDF diagram can be also seen in recent MD simulation on the solution containing N-methylacetamide (NMA) and trehalose [54]. Therefore trehalose works as not only hydrogen bonding donor but also hydrogen bonding acceptor towards protein backbone.

The presence of multiple hydroxyl groups on each ring of trehalose allows for the formation of a large number of inter-molecular hydrogen bonds [55]. Besides the hydrogen bonding interactions towards protein, trehalose can also form hydrogen bonds to other trehalose molecules as well as water molecules (e.g., see the high peak at ~2.78 Å in the RDF diagram of trehalose hydroxyl oxygens around water oxygen (Figure S5 in Supplementary Materials) referring to trehalose-water hydrogen bonding). To understand whether the strong hydrogen bonding ability of trehalose towards other species (protein and water) affects the hydrogen bonding interaction from water to protein, the RDF diagram of water oxygen around protein backbone carbonyl oxygen is presented in Figure 6 (a). A peak can be seen at ~2.70 Å, indicating the hydrogen bonding between protein backbone and water where the latter acts as a donor. Interestingly, the peak of RDF for protein in trehalose solution has similar height as that in pure water. In addition, the peak of the distribution of either length or angle of the hydrogen bonding between protein and water is located at same position and has similar height for both solutions (Figure 6 (b-c)). All of these results suggest that the formation manner and strength of protein-water hydrogen bonding interaction is not influenced by trehalose, consistent with the close electrostatic energy between individual water to protein in both solutions (Figure 5).

**Figure 6.**
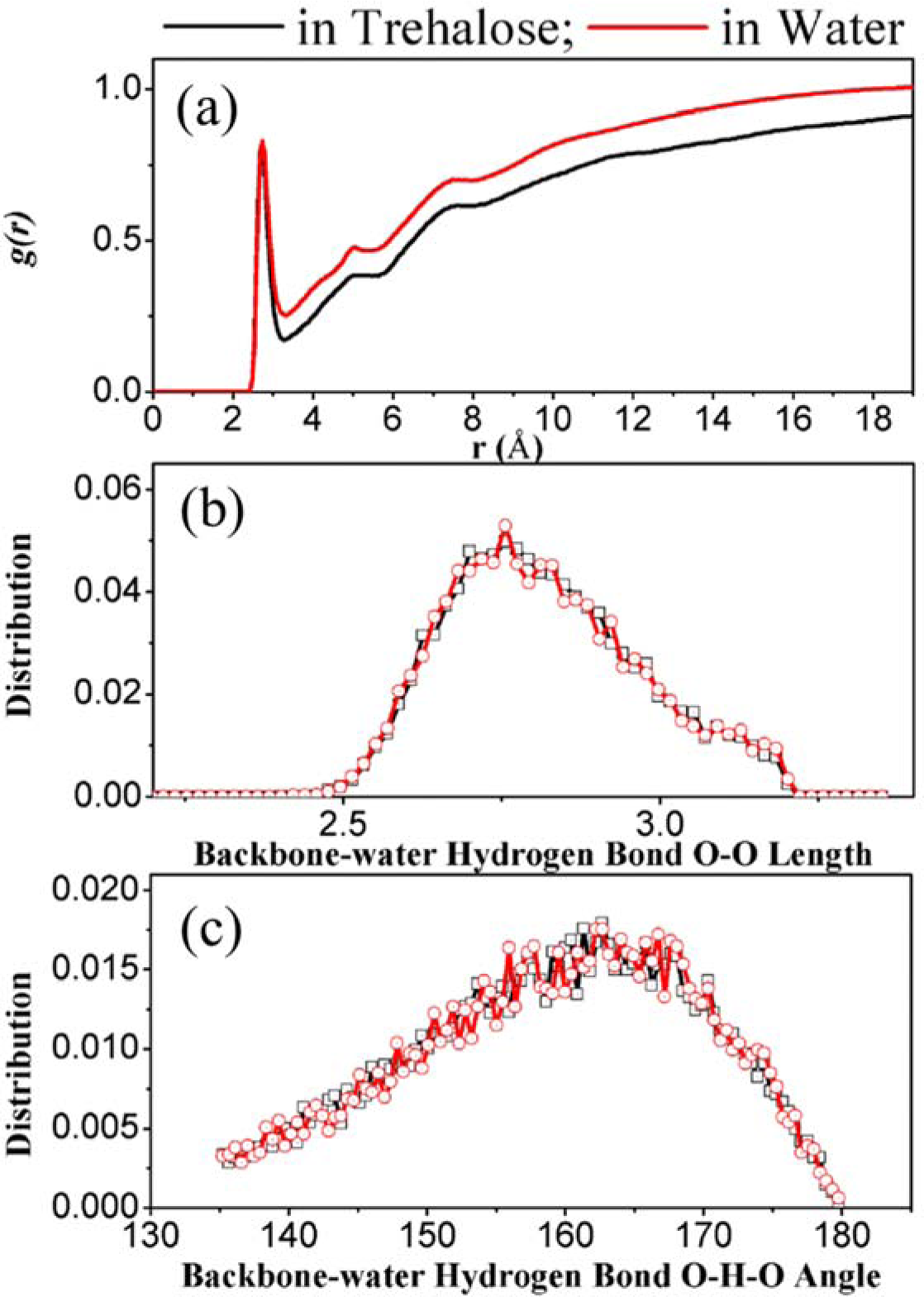
(a) Radial distribution function (RDF) of water oxygen around protein backbone carbonyl oxygen. (b) Length distribution and (c) angle distribution of hydrogen bonds between protein backbone carbonyl groups and water water molecules.

On the other hand, the RDF diagram of water oxygen around a central water oxygen has a high peak at ~2.80 Å and the height of the peak in trehalose solution is apparently larger than that in pure water (Figure 7 (a)). In addition, although the distribution of the hydrogen bonding angle among water molecules shares the same shape for both solutions, the distribution of water-water hydrogen bonding length is narrower around the center at ~2.80 Å in trehalose solution than that in pure water, implying that the presence of trehalose to some extent enhances the hydrogen bonding network among water.

**Figure 7.**
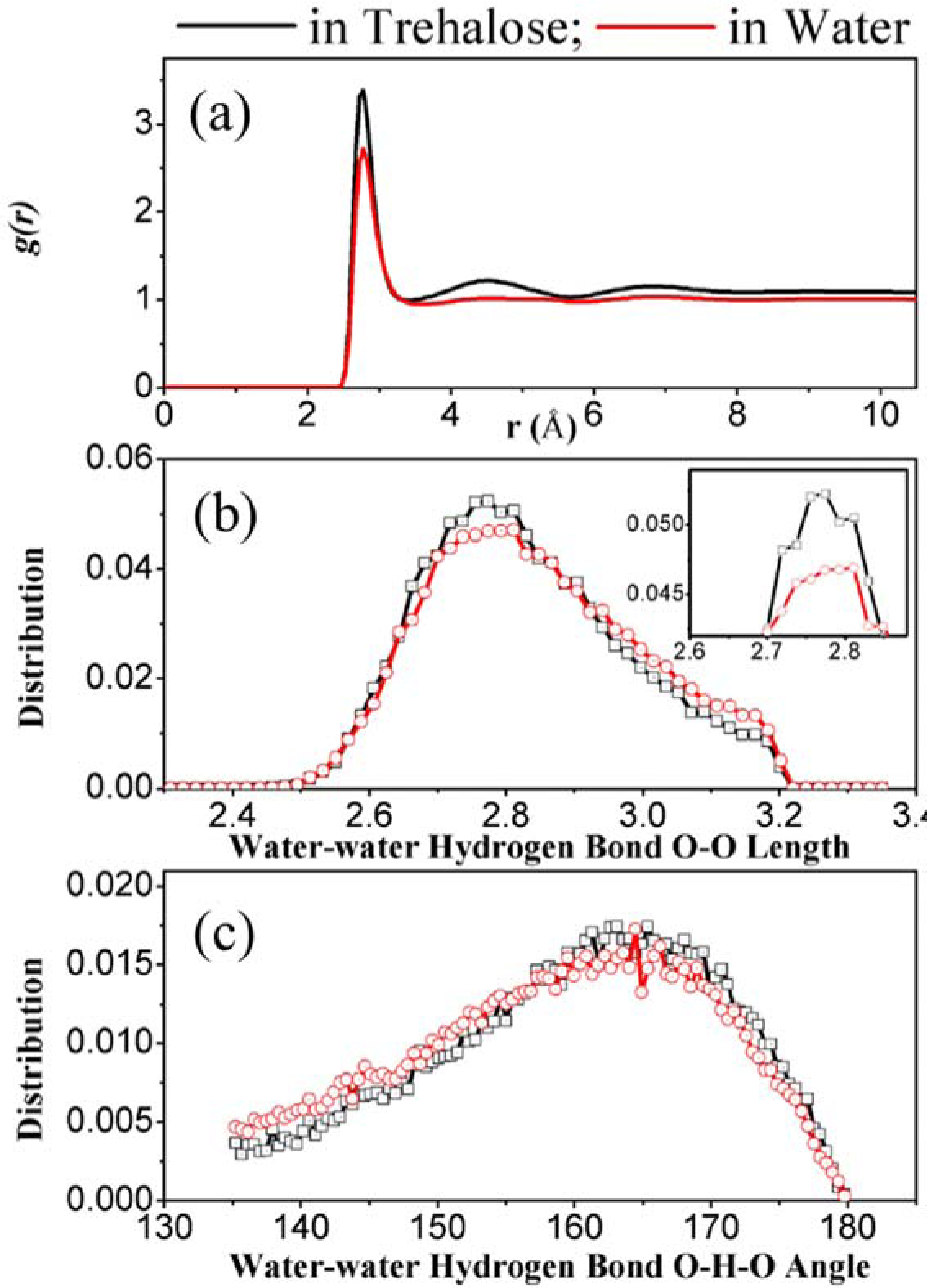
(a) Radial distribution function (RDF) of water oxygen around water oxygen. (b) Length distribution and (c) angle distribution of hydrogen bonds among water molecules.

### Trehalose Stabilizing Protein through Replacing Water to Form Hydrogen Bonds with Protein

Of intra-protein electrostatic interactions, the backbone hydrogen bonding between carbonyl and aminde groups is the main structural element and its stability is crucial for secondary structure. Water molecules, due to their small size, can insert between the carbonyl and amide groups and impair the intra-molecule hydrogen bonds [56]. Figure 8 shows the time series of the average numbers of backbone hydrogen bonds, and the hydrogen bonds from trehalose/water to protein backbone for the simulation of BBA5 in trehalose solution. Protein-water hydrogen bonds are the most among all kinds of hydrogen bonds, with the number being decreased from the beginning of simulation. Meanwhile, the protein-trehalose hydrogen bonds are arisen, with the increased number slightly smaller than the decreased number of protein-water hydrogen bonds. As shown in Figure 8 (b-c), the length of protein-trehalose hydrogen bonding is centered at ~2.73 Å, similar to the center of the length of protein-water hydrogen bonding (Figure 6 (b). The distribution of protein-trehalose hydrogen bonding angle is, however, broader and the center is slightly shifted to small angle range (blue-shifted), implying that the protein-trehalose hydrogen bonding might be not as strong as protein-water hydrogen bonding. As a result of the reduction of protein-water hydrogen bonds (and the increasing of protein-trehalose hydrogen bonds), the intra-protein backbone hydrogen bonds become stable, as revealed by the steady number of such hydrogen bonds during the entire simulation in Figure 8 (a). As a comparison, without the interruption of trehalose, the number of protein-water hydrogen bonds keeps large and as a result the number of intra-protein hydrogen bonds is relatively small for BBA5 in pure water (Figure S6 in Supplementary Materials).

**Figure 8.**
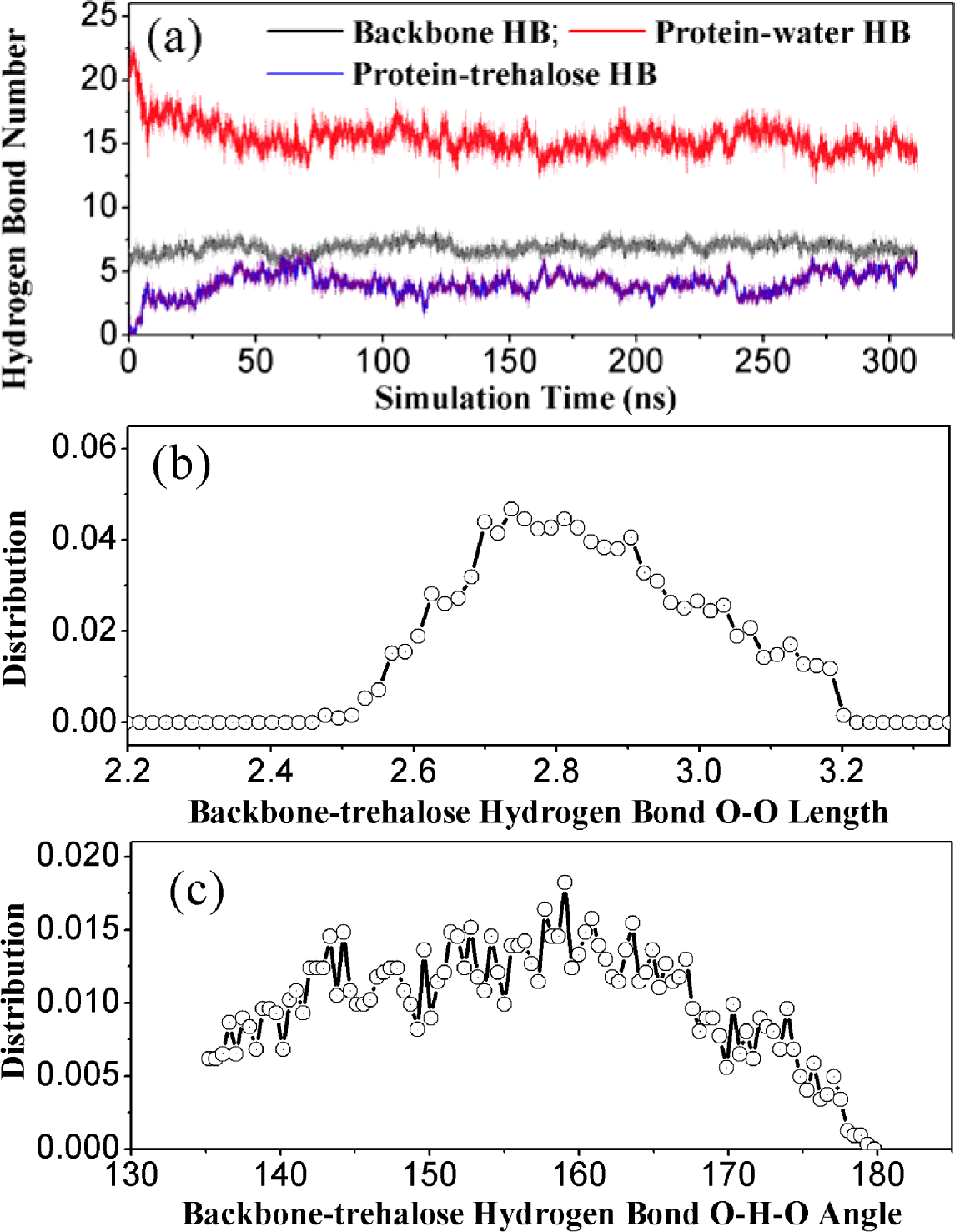
(a) Time series of intra-protein backbone hydrogen bonds as well as inter-protein hydrogen bonds from water and trehalose for BBA5 in trehalose solution. (b) Length distribution and (c) angle distribution of hydrogen bonds between protein backbone and trehalose molecules.

## Discussion and Conclusions

Using all-atom molecular dynamics simulation to measure the structural stability of a protein model (BBA5) in trehalose solution and pure water, the present comparative study attempted to investigate the molecular mechanism underlying the stabilizing effects of trehalose on protein from both structural and energetic perspectives. In principle, the accuracy of theoretical simulation depends on the choice of force field. The well-developed AMBER F99SB molecular force field [46] and TIP3P explicit solvent model [48] were used to model protein and water molecules, respectively. In addition, the parameters of the force field concerning trehalose were taken from GLYCAM06 force field [47], which has been widely used for molecular simulations of monosaccharides and oligosaccharides [57–59]. In comparison to other force fields such as CHARMM-based force field (CSFF), the GLYCAM06 force field could describe the structural and dynamical properties (e.g., self-diffusion constant) of several disaccharides more consistent with DFT calculations and experimental data [54,60,61]. In addition, a comparison of different disaccharide force fields (GLYCAM06, PM3-CARB1, and SCC-DFTB-D) indicated that when combined with TIP3P water model, the GLYCAM06 force field is the best at predicting experimentally consistent three-dimensional conformation of monosaccharide puchering and solvent interactions [62].

The present molecular simulation indicates that trehalose has favorable electrostatic interaction energy in aqueous solution owing to the large amount of hydrogen bonding agents within trehalose. As a result, trehalose molecules have strong tendency to cluster with each other through inter-molecular hydrogen bonding interactions. In addition, hydrogen bonding interactions can be also formed among trehalose and water. The involvement of trehalose in the hydrogen bonding network in bulk solution to some extent influences the hydrogen bonding strength among water molecules (Figure 7), but such influence can be not seen for the hydrogen bonding from water to protein (Figure 6). On the other hand, trehalose also has a moderate tendency to approach protein of which two events are involved: 1) trehalose can form hydrogen bonds with protein; 2) water molecules are partially expelled from protein surface and the hydrogen bonds between protein and water are thus reduced. Therefore, the protein-water hydrogen bonding interactions are replaced by (weaker) protein-trehalose hydrogen bonding, which induces the enhancement of protein structure stability in trehalose solution. This observation is in accordance with the “water replacement hypothesis” proposed in previous research study [3,20,21]. In summary, the present study provides a picture of the molecular interactions in protein-trehalose-water complex system and displays that the water replacement scenario is adequate to interpret trehalose-induced protein stabilization in low-concentrated trehalose solution.

## Acknowledgements

This work was supported by research grants from the National Natural Science Foundation of China (Grand No. 21373258 and 21003003) and National 863 Program (Grant No. 2012AA01A305). For the computations in this article, we used the computation resources in Shanghai Supercomputer Center (SSC), and TianHe-1 supercomputer in Tianjin.

