## Supplementary Materials for "Trehalose Stabilizing Protein in a Water Replacement Scenario: Insights from Molecular Dynamics Simulation"

| Solution | Trehalose | H_2_O | Simulation Length | |
| --- | --- | --- | --- | --- |
| Trehalose/water | 120 | 6949 | Traj 1 | 340 ns |
|  |  |  | Traj 2 | 338 ns |
|  |  |  | Traj 3 | 312 ns |
| Water | --- | 5639 | Traj 1 | 315 ns |
|  |  |  | Traj 2 | 324 ns |
|  |  |  | Traj 3 | 314 ns |

Table S1. Parameters for individual simulations.

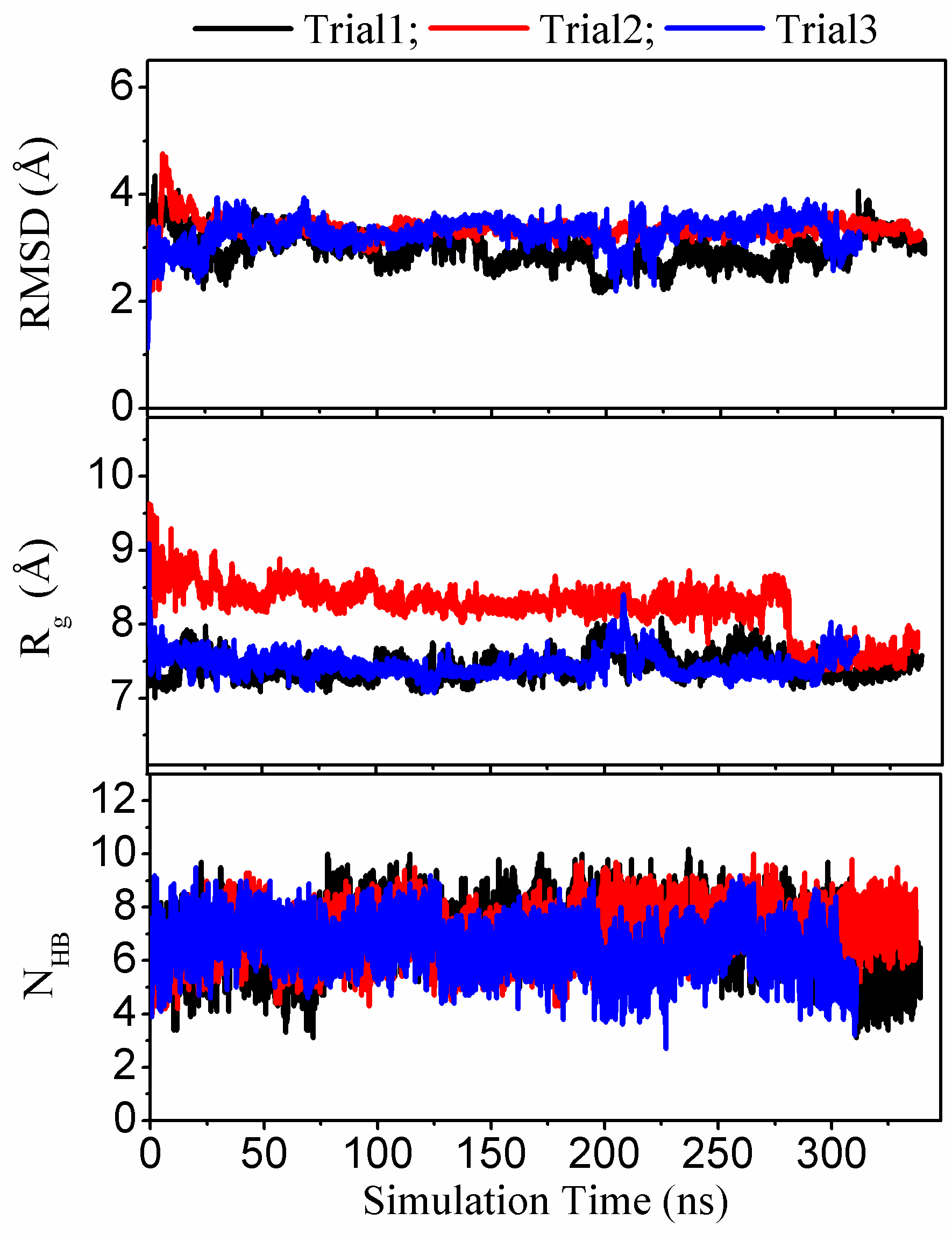

Figure S1. Time series of (a) the root-mean-square deviation (RMSD) corresponding to the NMR structure, (b) the radius of gyration (*R_g_*), and (c) the total number of backbone hydrogen bonds formed within BBA5 protein from individual trajectories of trehalose aqueous solution respectively.

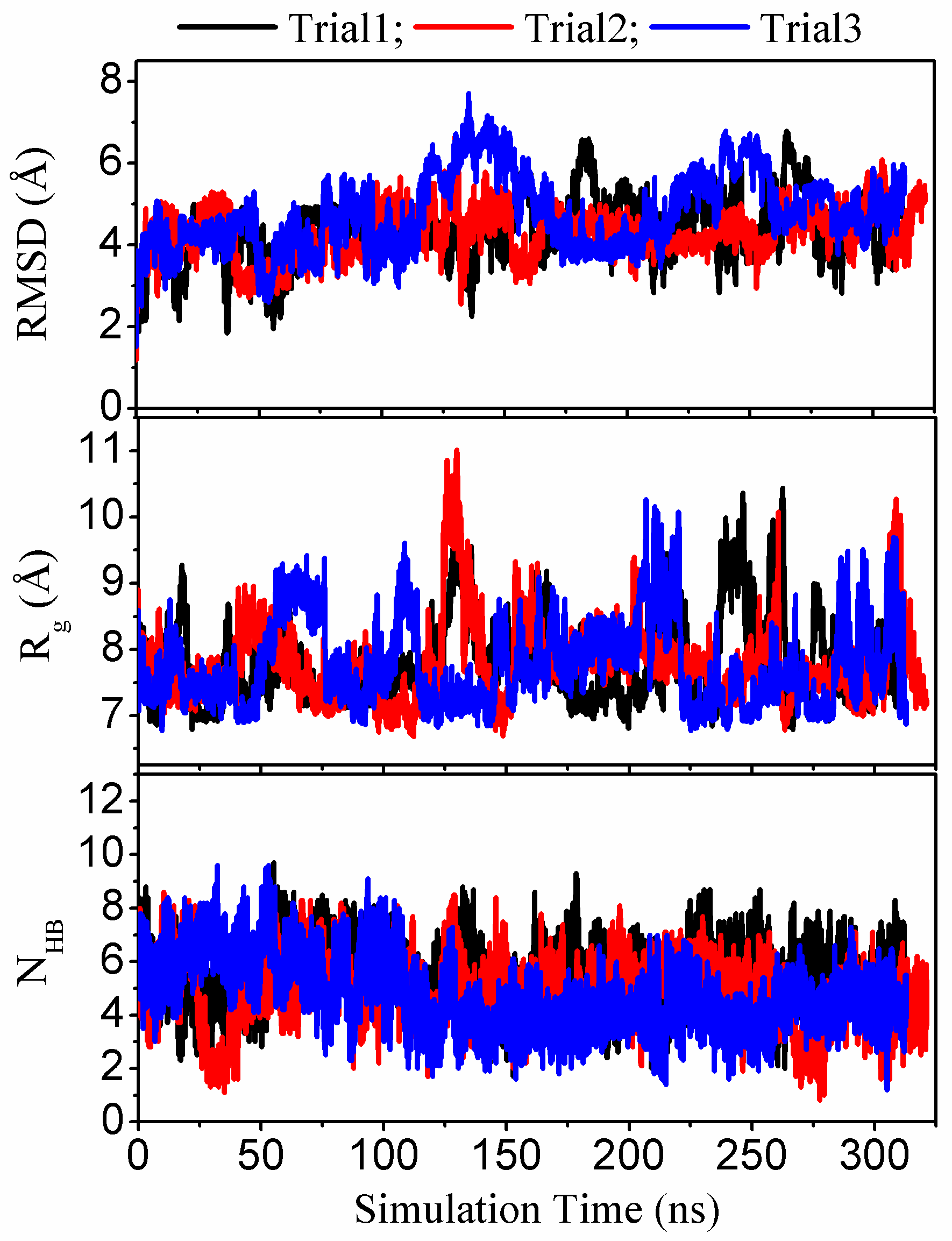

Figure S2. Time series of (a) the root-mean-square deviation (RMSD) corresponding to the NMR structure, (b) the radius of gyration (*R_g_*), and (c) the total number of backbone hydrogen bonds formed within BBA5 protein from individual trajectories of pure water respectively.

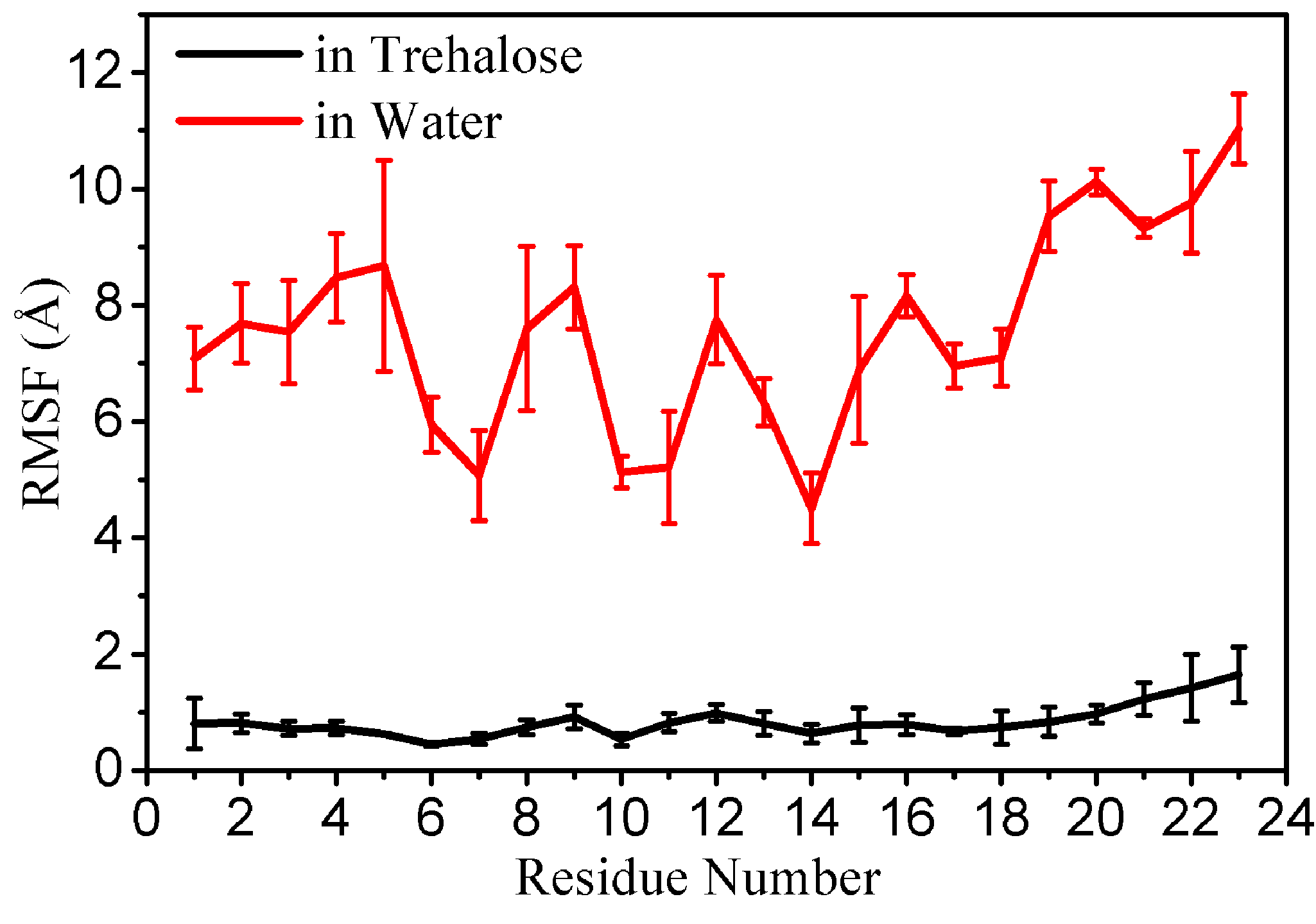

Figure S3. Root-mean-square-fluctuation (RMSF) *vs* sequence index for BBA5 protein in trehalose solution and pure water, respectively. The parameter value is calculated covering all trajectories for each simulation system and the data from single trajectories is used to calculate error bars.

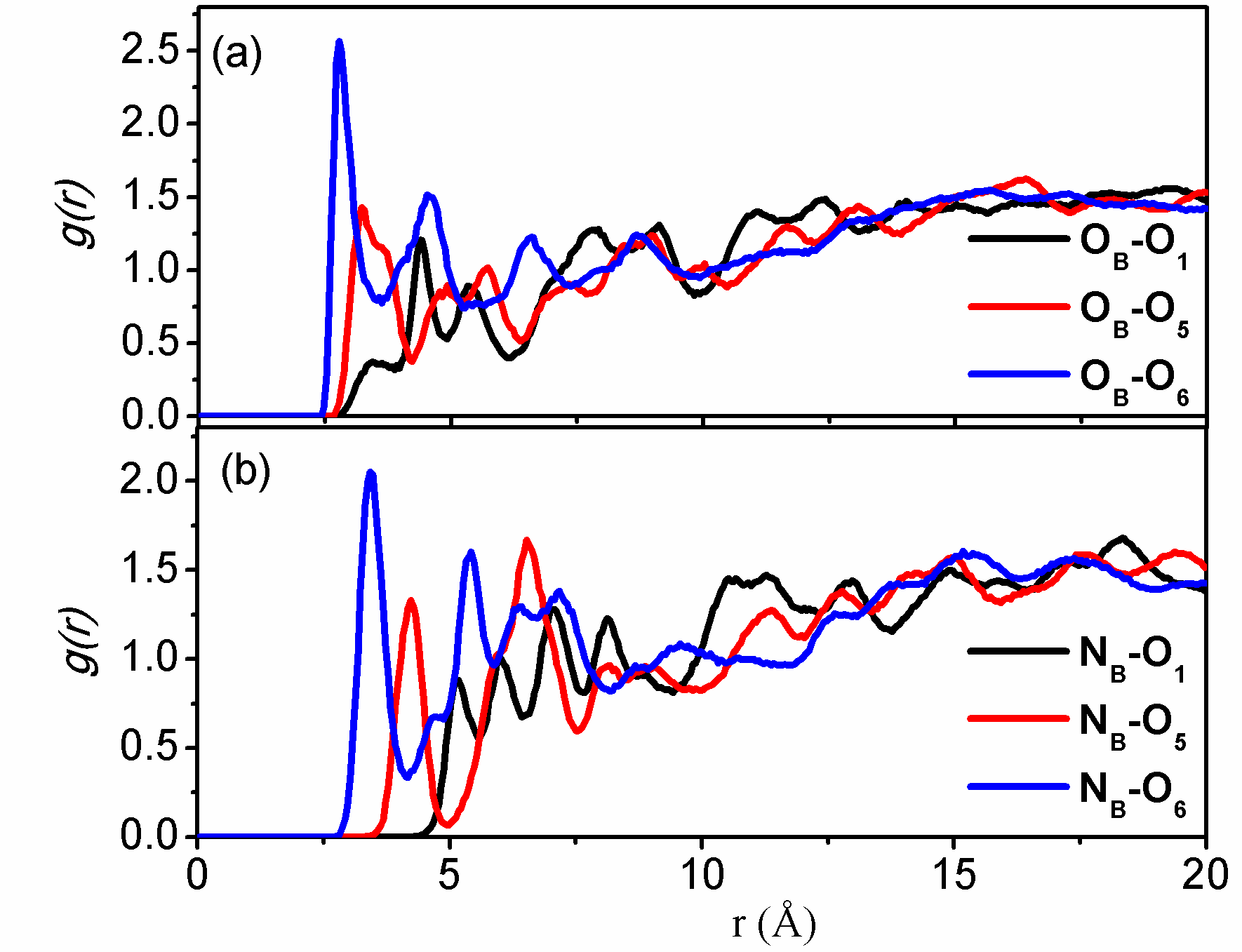

Figure S4. Site-site radial distribution functions between backbone carbonyl/amide groups and trehalose hydroxyl oxygens (O_1_, O_5_, O_6_) for BBA5 in trehalose solution. O_B_: backbone carbonyl oxygen; N_B_: backbone amide nitrogen.

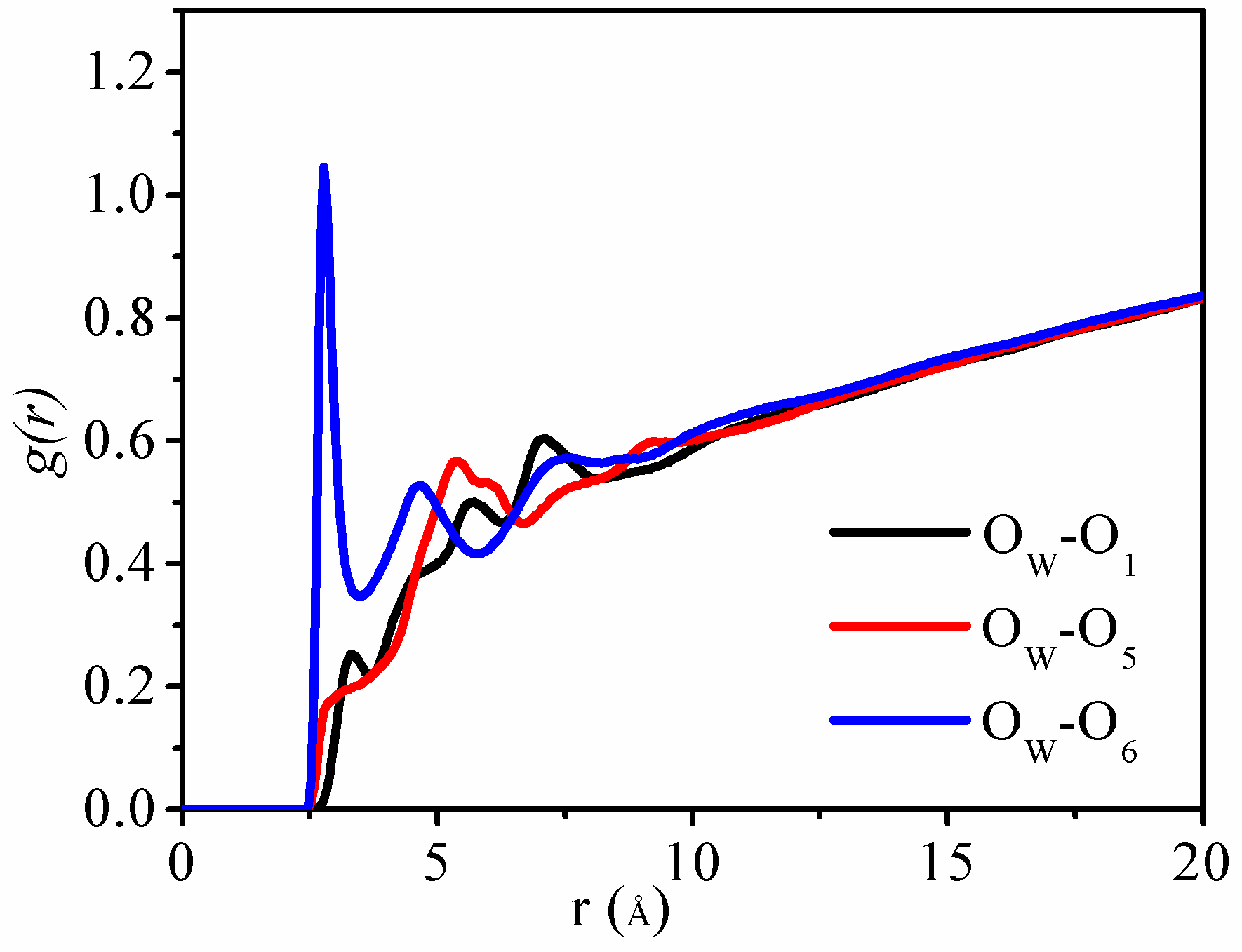

Figure S5. Site-site radial distribution functions between trehalose hydroxyl oxygens (O_1_, O_5_, O_6_) and water oxygen atom (O_W_) for BBA5 in trehalose solution.

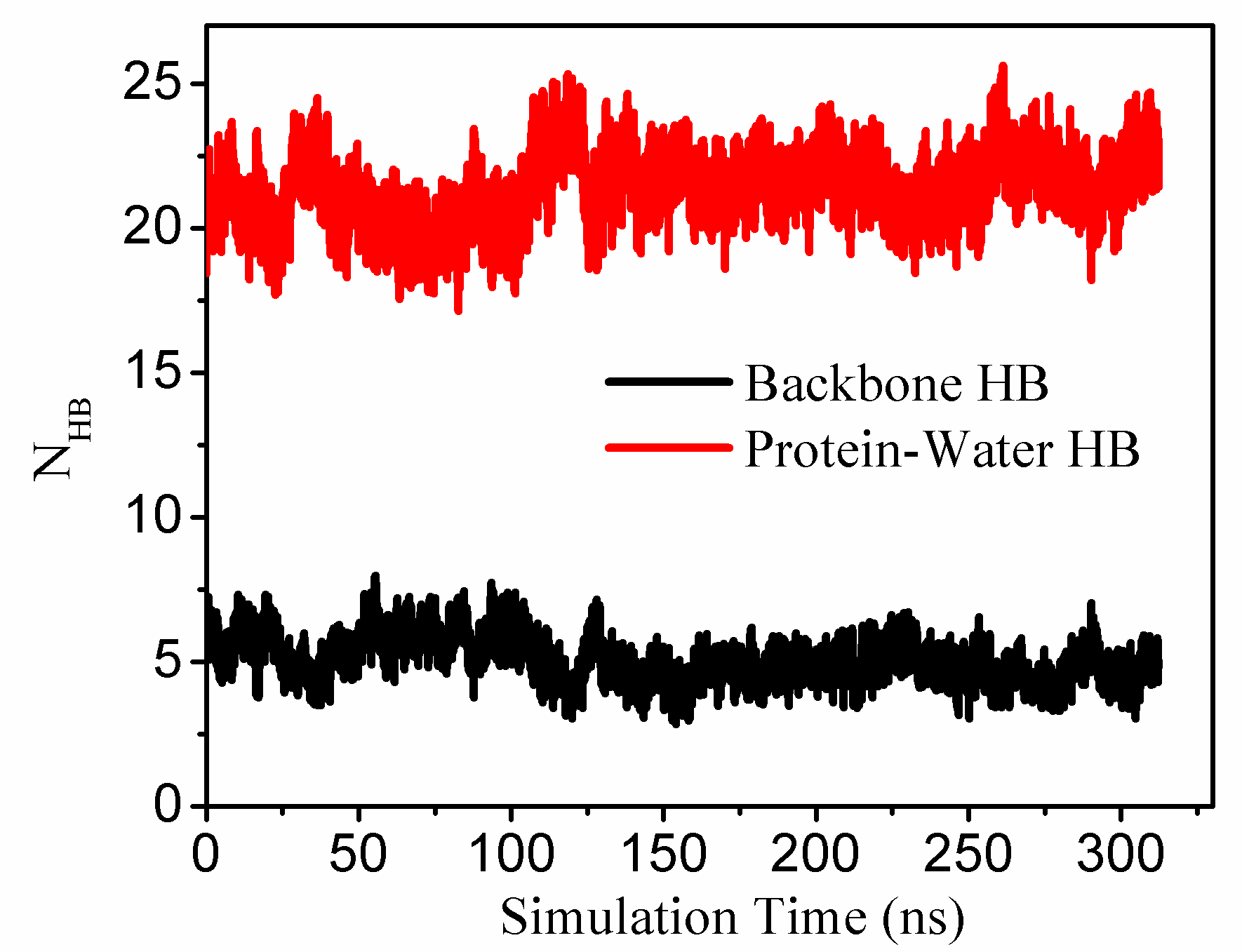

Figure S6. Time series of intra-protein backbone hydrogen bonds and inter-protein hydrogen bonds from water for BBA5 in pure solution.
